# Carbonyl Post-Translational Modification Associated with Early Onset Type 1 Diabetes Autoimmunity

**DOI:** 10.1101/2021.06.15.448522

**Authors:** Mei-Ling Yang, Sean E. Connolly, Renelle Gee, TuKiet Lam, Jean Kanyo, Steven G. Clarke, Catherine F. Clarke, Eddie A. James, Cate Speake, Carmella Evans-Molina, Li Wen, Kevan C. Herold, Mark J. Mamula

## Abstract

Inflammation and oxidative stress in pancreatic islets amplify the appearance of various post-translational modifications (PTMs) to self-proteins. Herein, we identified a select group of carbonylated islet proteins arising before the onset of hyperglycemia in non-obese diabetic (NOD) mice. Of particular interest, we identified carbonyl modification of the prolyl-4-hydroxylase beta subunit (P4Hb) that is responsible for proinsulin folding and trafficking as an autoantigen in both human and murine type 1 diabetes. We found the carbonylated-P4Hb is amplified in stressed islets coincident with decreased glucose-stimulated insulin secretion and altered proinsulin to insulin ratios. Moreover, circulating autoantibodies against P4Hb were detected in prediabetic NOD mice and in early human type 1 diabetes prior to the onset of anti-insulin autoimmunity. Our studies provide mechanistic insight into the pathways of proinsulin metabolism and those creating autoantigenic forms of insulin in type 1 diabetes.

## INTRODUCTION

Type 1 diabetes is an organ-specific autoimmune disease marked by lymphocyte infiltration of pancreatic islets. Insulitis results in the liberation of reactive oxygen species (ROS) and induces the production of proinflammatory cytokines. Accumulating evidence in both experimental and clinical studies demonstrate that oxidative stress induced by hyperglycemia and insulitis plays a key role of the onset of type 1 diabetes and diabetes-related complications of disease (1; 2). For example, islet proteins exposed to metal-catalyzed oxidation generate glutamic acid decarboxylase (GAD65) aggregates that react with serum autoantibodies from type 1 diabetes patients (3). Moreover, antioxidants delay and/or prevent the onset of type 1 diabetes in non-obese diabetic (NOD) and streptozotocin (STZ)-induced type 1 diabetes murine models (4; 5). Of interest, oxidative stress usually accompanies a wide spectrum of post-translational protein modifications (PTMs) such as carbonylation, hydroxylation, nitrosylation and glutathionylation (6). PTMs are known as one mechanism by which autoreactive T and B cells escape selection and break tolerance to neo self-proteins (1). In the present study, we examine protein modifications induced by inflammation and oxidative stress leading to neo-autoantigens in type 1 diabetes.

Protein carbonylation is a major product of tissue proteins in response to oxidative stress. Both oxidative stress and protein carbonylation contribute to insulin resistance and metabolic dysfunctions in adipose tissue of both animal models and human type 1 diabetes (7). Carbonylation leads to sarco(endo)plasmic reticulum Ca^2+^-ATPase (SERCA2a) activity loss and diastolic dysfunction in the STZ-induced type 1 diabetes murine model (8). Recent studies profiled carbonylated plasma proteins as potential biomarkers in type 2 diabetes (T2D) (6; 9). Little is known about the role of carbonylated neoantigens in the progression of type 1 diabetes.

Herein, proteomic analysis identified 23 carbonylated islet polypeptides unique to pre-diabetic NOD mice. Seven of these polypeptides were recognized by serum from pre-diabetic NOD mice, including protein disulfide isomerase (PDI) isoforms, 14-3-3 protein isoforms, glucose-regulated protein 78 (GRP78) and chymotrypsinogen B. Of interest, prolyl-4-hydroxylase beta (P4Hb, also known as protein disulfide isomerase A1; PDIA1) was found to be an early autoantigen in both human and murine models of type 1 diabetes. P4Hb is required for proinsulin maturation and pancreatic beta cell health (10). Our data suggest a novel role of modified P4Hb, both as an early target of autoimmunity as well as a pathway that provokes autoimmunity to insulin and/or proinsulin. As discussed herein, modified P4Hb biological functions provide explanation for recent observations of increased proinsulin to insulin ratios in the progression of type 1 diabetes (11).

## RESEARCH DESIGN AND METHODS

### Animals and type 1 diabetes serum samples

NOD/ShiLt, BALB/c, and C57BL/6 mice were purchased at 3 wks of age from Jackson Laboratories (Bar Harbor, Maine). Experiments were performed in female mice unless indicated for male mice. All mice were housed in microisolator cages with free access to food and water and maintained at the Yale Animal Resource Center at Yale University. All animal studies were performed in accordance with the guidelines of the Yale University Institutional Animal Care and Use Committee. Glucose levels were measured in blood withdrawn from the retroorbital sinus of anesthetized mice using Lite glucometer and test strips (Abbott Laboratories, Abbott Park, IL, USA). Mice with blood glucose values greater than 250 mg/dL were considered diabetic. Serial serum samples and lymph node cells were acquired from NOD and age matched control animals. Serum glucose was measured as indicated. Human samples were collected under from subjects with healthy control subjects (n=10), established type 1 diabetes (n=12, C peptide<0.3pmol/ml, from 5-40 years after type 1 diabetes diagnosis) and early onset disease (n=21, C peptide >0.4pmol/ml), at different time point before and/or after insulin treatment (1, 3, 6, 9, and 12 months). IRB approval for the study was sought and received from the Benaroya Research Institute IRB. Samples were assayed in a blinded fashion for binding to P4Hb as described below. C-peptide levels were measured at Northwest Lipid Research Laboratories using a two-site immunoenzymatic assay on the Tosoh II 600 autoanalyzer.

### Pancreatic islet isolation/culture and protein extraction

For murine samples, pancreatic islets were hand-picked from 5-week old NOD mice, as previously described (12). After three freeze-thaw cycles, the islet protein concentration was measured by a modified Lowry assay as described previously.

For human samples, the pancreatic islets were obtained from deceased organ donors provided by the cGMP Cell Processing Facility at the University of Miami Miller School of Medicine (Miami, Florida) in accordance with ethical regulations. All donors (n=6) were 25-53 years old with no history of diabetes or other pancreatic disease. The purity and viability of islets were > 90% as determined by dithizone staining. Islet aliquots (2,000 islet equivalent; IEQ) were cultured in CMRL1066 culture medium (Corning) containing 10% FCS, 2 mM L-glutamine, and 100 U/ml penicillin/streptomycin at 37°C in 5% CO_2_ humidified incubator. For oxidative or inflammatory stress experiments, human islets were incubated with either 500µM H_2_O_2_ for 1 h or rhINFγ (1000U/ml), rhIL-1β (50U/ml) and rhTNFα (1000U/ml) for 68-72 h. For glucose-stimulated insulin release assay, human islets were washed free of H_2_O_2_ or cytokines and treated with either 5.5mM or 16.7mM D-(+)-glucose (Sigma) for 1 h. Supernatants were then assayed for insulin and proinsulin concentration by commercial ELISA (ALPCO, Inc. Salem, NH).

### Differential 2D gel electrophoresis

Islet protein extracts were separated by standard 2D-gel electrophoresis by using ZOOM IPGRunner first-dimension isoelectric focusing unit (Invitrogen) and followed by second-dimension 12.5% SDS-PAGE. Prior to isoelectric focusing (IEF), proteins were precipitated away from detergents, salts, lipids, and nucleic acids using the 2-D Clean-Up Kit (GE Healthcare, Pittsburgh, PA). Samples were denatured for 60 min at room temperature in sample buffer containing ZOOM 2D protein solubilizer I, 10 mM DTT, and 0.5% (w/v) 3-10 carrier ampholytes (Invitrogen). Rehydration of ZOOM Strip (pH 3-10, non-linear, Invitrogen) was carried out overnight at room temperature. After IEF, strips were alkylated with 125 mM iodoacetamide (Sigma). Second dimension resolution was carried out by standard SDS-PAGE, followed by protein staining or transfer to nitrocellulose for immunoblotting. Silver staining of 2D gels prior to spot excision and mass-spectroscopy was carried out using the SilverQuest silver staining kit (Invitrogen).

### Detection of carbonyl modification by OxyBlot

Protein carbonylation from islet lysates was detected by using OxyBlot^TM^ Protein Oxidation Detection Kit (Millipore). Briefly, carbonyl groups were derivatized by 2,4 dinitrophenylhydrazine (DNPH) to form a stable dinitrophenyl (DNP) hydrozone product and then detected by anti-DNP antibodies. Bands were visualized by chemiluminescent detection on HyBlot CL autoradiography film (Denville Scientific).

### Identification of the carbonyl modification of P4Hb by DNPH-assisted mass spectrometry

The purified rhP4Hb was incubated in PBS (native) or PBS containing 100µM FeSO4, ascorbate, 25mM H_2_O_2_ and 25mM ascorbate (oxidation) at 37°C for 4h and kept at −80°C in multiple aliquots to avoid freeze and thaw until used (13). The native and oxidative rhP4Hb were derivatized with DNPH as previously described with some modifications (14). In brief, rhP4Hb (100µg) was incubated with 5mM DNPH in PBS for 30 min at room temperature, neutralized by 0.5mM Tris base and washed out excess DNPH and concentrated in PBS by Amicon Ultra centrifugal filter (5K). Then DNPH-derivatized rhP4Hb was trypsinized for MS/MS analysis and re-derivatized with DNPH as described above. After the second time of DNPH derivatization, the trypsinized DNPH-derivatized rhP4Hb peptides were incubated in the presence or absence of 15 mM sodium cyanoborohydride (stabilizer), which can stabilize the hydrazine bond between the hydrazide and the carbonyl group, for 30 min on ice with occasional shaking for following MS/MS analysis. In the absence of stabilizer, the mass shifts used for identification of DNPH-derivatized carbonyl group of arginine, proline, lysine, and threonine, were 136.97, 194.11, 179.02 and 178.01, respectively (15). With stabilizer, the mass shifts were added on by 2 daltons for each residue (16).

### Mass spectrometry and data searching

LC MS/MS analysis of unknown proteins was performed on a Waters Q-Tof Ultima mass spectrometer at the W.M. Keck Facility at Yale University. All MS/MS spectra were searched using Thermo Proteome Discoverer software (version 2.2.0.388) linked to the automated MASCOT algorithm (version 2.7.0) against the NCBI nr database. Then MS/MS spectra and carbonylation site were validated by Scaffold (version 4.8.9) and Scaffold PTM (version 4.0.1).

### Lymphocyte Proliferation Assay

Pooled lymph node cells including pancreatic lymphoid cells from age-matched mice were isolated and followed by traditional [^3^H] thymidine incorporation proliferation assay as described previously (17). In brief, cells (5 x 10^5^) were plated in 96-well flat-bottom microtiter plates with pre-coated anti-CD3 (10µg/ml) mAb as positive control or different concentrations of rhP4Hb protein. Cells were incubated for 48 h after which cells were pulsed with 1 µCi [^3^H] thymidine (ICN Chemicals, Irvine, CA) for 18 h before being harvested onto filters with a semiautomatic cell harvester. Radioactivity was counted with a Betaplate liquid scintillation counter (Perkin Elmer Wallac). Recombinant human P4Hb protein was expressed and purified from a bacterial expression vector (hPDI-pTrcHisA; a generous gift from Professor Colin Thorpe, University of Delaware, Newark, Delaware), as previously described (18).

### Detection of carbonylated-P4Hb and oxidative status by flow cytometry and confocal microscopy

For flow cytometry, human islets were dissociated into single-cell suspension by accutase (Millipore) and the islet cells were fixed by 95% methanol for 30 min on ice. Islet cells were stained with anti-P4Hb (Abcam) and goat-anti-rabbit Alexa Fluor 647 (Life Technology) in staining buffer (PBS containing 1% BSA and 0.1% Tween-20). Islet cells were then treated with 1mM DNPH (Fluka) or 2N HCl (as control derivatization buffer) in the above staining buffer containing 0.1% SDS for 15 min in dark at room temperature. Cells were stained with anti-DNP (Sigma) at 4 °C overnight and followed by incubation with secondary antibody, anti-goat Alexa Fluor 488 (Abcam). Intracellular ROS level was measured by CellROX*^®^* Deep Red probe (5 µM, 30 min at 37 °C, Molecular probes) according to the manufacturer’s instructions. Stained cells were analyzed by FACSCalibur (BD Biosciences) with FlowJo software (Tree Star).

For confocal microscopy, the 4 chamber glass slide (Falcon) was pre-treated with 1% Alcian blue solution for 15 min at room temperature and washed four times with H_2_O. The intact human islet suspension was applied to Alcian Blue-coated slides and centrifuge at 300rpm for 3 min. The islets were fixed with 2% paraformaldehyde in PBS for 30 min, washed twice with PBS and permeabilized with 0.3% Triton X-100 for 2 h. The islets were treated with DNPH or HCl in staining buffer (PBS, 0.5%BSA, 0.05% Tween-20) containing 0.1% SDS. After derivatization, the islets were stained with anti-insulin (R&D Systems), anti-P4Hb and anti-DNP, respectively, as described above. Islets were then observed on a Leica SP5 II laser scanning confocal microscope.

### ELISA and immunoblot assay

Immunoreactivity of human and mouse serum to P4Hb and insulin was performed by ELISA. Briefly, 1 µg of recombinant human P4Hb protein or insulin (Gemini Biotech) in 0.05 M carbonate-bicarbonate buffer (pH = 9.6; Sigma) was coated onto ELISA plates (Thermo Scientific) overnight at 4°C and blocked with 1%BSA in PBST or commercial Block^TM^ Casein (Pierce Biotechnology, Rockford, IL) containing 1% Tween-20. Sera were diluted as 1:100 in diluting buffer (0.03% BSA in PBST) and incubated 2 h at room temperature. Species-specific goat anti-IgG or anti-IgM alkaline phosphatase was used as a secondary reagent (Southern Biotech). Color was developed via the addition of pNPP substrate (Sigma) and absorbance was read at 405nm (Synergy HT Multi-Mode Reader, BioTek Instruments). Individual signals were normalized to no-antigen control wells. Mouse polyclonal antibody against P4Hb protein (Abcam) served as positive control in ELISA. All readings were normalized to non-specific serum binding to no-antigen control wells. Autoantibodies against to P4Hb or insulin was designating as positive with an OD > 2 standard deviation (SD) above BALB/c serum or human healthy serum.

For immunoblotting, protein samples (pancreatic islet extract isolated from NOD mice) was separated by 2D-SDS-PAGE, blotted onto a nitrocellulose membrane, and probed with serum (1:100) from NOD mice or from type 1 diabetes patients and incubated with the alkaline phosphatase-conjugated anti-mouse or anti-human IgG, then visualized by NBT/BCIP substrate (Thermo Scientific).

### Statistical analysis

All statistics were performed using a Student’s unpaired two-tailed *t* test. A value of *p* < 0.05 was regarded as significant.

## RESULTS

### NOD islet proteomic analyses for carbonyl-modified proteins

Pancreatic islet extracts were prepared from NOD and NOD-SCID animals and examined by gel electrophoresis. Protein carbonylation was visualized by derivatization with 2,4-dinitrophenylhydrazine (DNPH) and probed with anti-DNP antibody. Not surprisingly, we found that prediabetic and diabetic NOD mice as well as NOD-SCID islet cell proteins had both similar and divergent patterns of carbonyl posttranslational modifications (data not shown). With particular interests in identifying carbonyl modification in pancreatic islets prior to disease onset, pancreatic islets proteins from 5-week-old prediabetic NOD mice were investigated by two-dimensional (2D)-gel electrophoresis and probed with anti-DNP antibody. Proteins from a total of 7 identified spots (as indicated in supplementary Fig. 1) were analyzed by LC MS/MS mass spectrometry and MASCOT database search. A number of both novel and established disease derived autoantigens were identified with carbonyl modifications (Table 1). The list includes protein disulfide isomerase A1 (PDIA1; P4Hb), PDIA2 and Hspa5 78kDa glucose related HSP (also known as GRP78, BiP), all abundantly carbonylated from prediabetic NOD islets (Table 1). Of interest, pancreatic PDIA2 is an autoantigen in CTLA-4 deficient mice characterized by a fatal lymphoproliferative disorder (19). Moreover, citrullinated GRP78 has been identified as an autoantigen in type 1 diabetes(20; 21). Two carbonyl-modified proteins identified from prediabetic NOD islet preparations, pancreatic amylase and chymotrypsinogen, were previously identified as biomarkers for autoimmune pancreatitis and fulminant type 1 diabetes (22; 23), respectively.

**Table 1.**
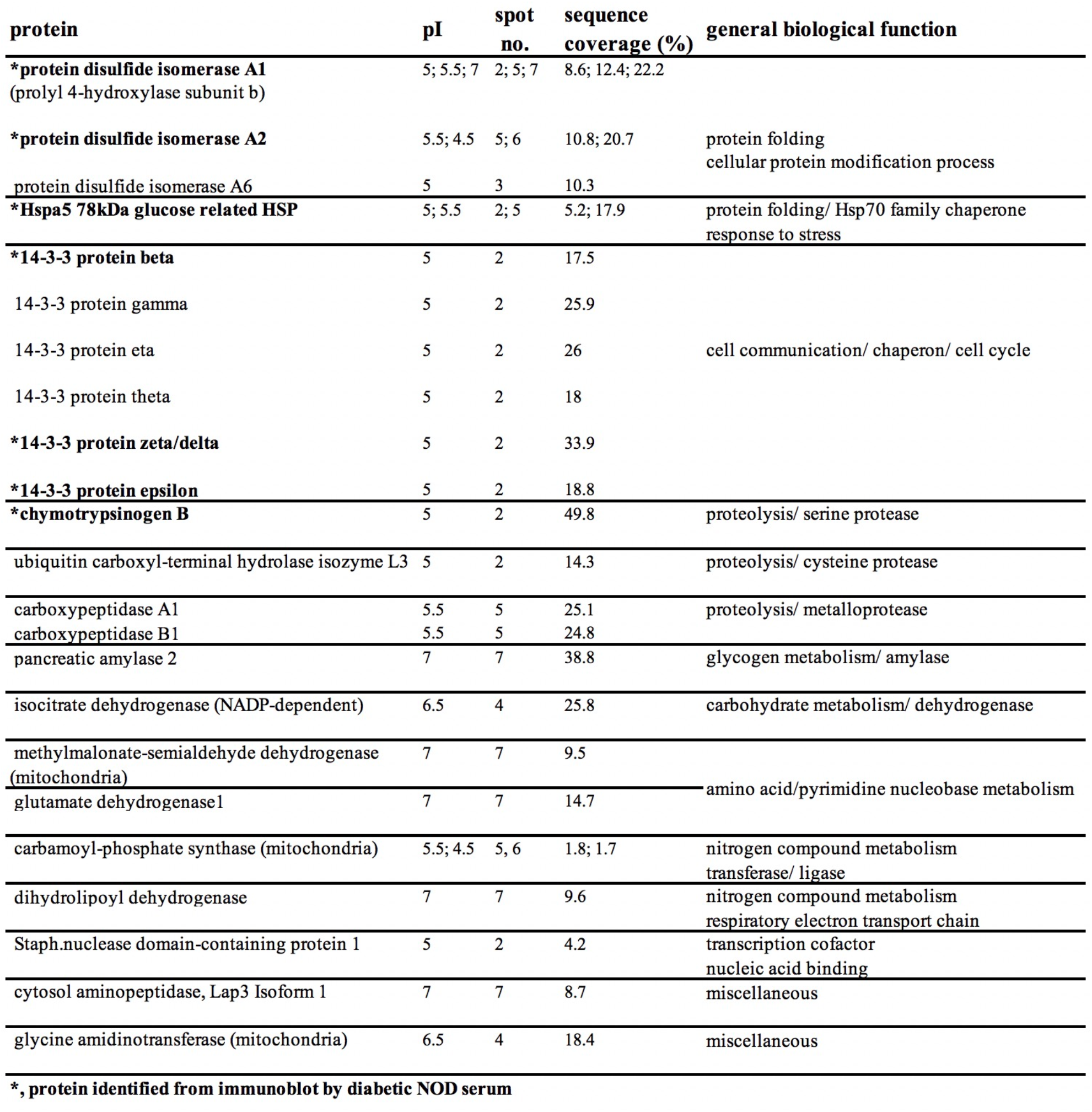
Identification of carbonylated islet proteins in pre-diabetic NOD mice.

| **protein** | **pI** | **spot no.** | **sequence coverage (%)** | **general biological function** |
| --- | --- | --- | --- | --- |
| **\*protein disulfide isomerase A1**<br>(prolyl 4-hydroxylase subunit b) | 5; 5.5; 7 | 2; 5; 7 | 8.6; 12.4; 22.2 |  |
| **\*protein disulfide isomerase A2** | 5.5; 4.5 | 5; 6 | 10.8; 20.7 | protein folding<br>cellular protein modification process |
| protein disulfide isomerase A6 | 5 | 3 | 10.3 |  |
| **\*Hspa5 78kDa glucose related HSP** | 5; 5.5 | 2; 5 | 5.2; 17.9 | protein folding/ Hsp70 family chaperone<br>response to stress |
| **\*14-3-3 protein beta** | 5 | 2 | 17.5 |  |
| 14-3-3 protein gamma | 5 | 2 | 25.9 |  |
| 14-3-3 protein eta | 5 | 2 | 26 | cell communication/ chaperon/ cell cycle |
| 14-3-3 protein theta | 5 | 2 | 18 |  |
| **\*14-3-3 protein zeta/delta** | 5 | 2 | 33.9 |  |
| **\*14-3-3 protein epsilon** | 5 | 2 | 18.8 |  |
| **\*chymotrypsinogen B** | 5 | 2 | 49.8 | proteolysis/ serine protease |
| ubiquitin carboxyl-terminal hydrolase isozyme L3 | 5 | 2 | 14.3 | proteolysis/ cysteine protease |
| carboxypeptidase A1 | 5.5 | 5 | 25.1 | proteolysis/ metalloprotease |
| carboxypeptidase B1 | 5.5 | 5 | 24.8 |  |
| pancreatic amylase 2 | 7 | 7 | 38.8 | glycogen metabolism/ amylase |
| isocitrate dehydrogenase (NADP-dependent) | 6.5 | 4 | 25.8 | carbohydrate metabolism/ dehydrogenase |
| methylmalonate-semialdehyde dehydrogenase<br>(mitochondria) | 7 | 7 | 9.5 |  |
| glutamate dehydrogenase1 | 7 | 7 | 14.7 | amino acid/pyrimidine nucleobase metabolism |
| carbamoyl-phosphate synthase (mitochondria) | 5.5; 4.5 | 5, 6 | 1.8; 1.7 | nitrogen compound metabolism<br>transferase/ ligase |
| dihydrolipoyl dehydrogenase | 7 | 7 | 9.6 | nitrogen compound metabolism<br>respiratory electron transport chain |
| Staph.nuclease domain-containing protein 1 | 5 | 2 | 4.2 | transcription cofactor<br>nucleic acid binding |
| cytosol aminopeptidase, Lap3 Isoform 1 | 7 | 7 | 8.7 | miscellaneous |
| glycine amidinotransferase (mitochondria) | 6.5 | 4 | 18.4 | miscellaneous |

**\*, protein identified from immunoblot by diabetic NOD serum**

### Identification of P4Hb as an antigenic islet protein

Thereafter, 2D gel electrophoresis of islet lysates from prediabetic NOD mice were next immunoblotted with selected diabetic NOD serum (Fig. 1). Protein spots bound by NOD serum were identified via LC MS/MS following by MASCOT database analysis. By this approach, P4Hb (also known as PDIA1) was identified as one major target of autoantibodies as indicated in Fig. 1. Other major carbonyl modified target proteins identified in immunoscreen by using diabetic NOD serum (Fig. 1) and in OxyBlot analysis from prediabetic NOD islet proteins (supplementary Fig. 1) were PDIA2, GRP78, 14-3-3 isoforms and chymotrypsinogen B (highlighted with asterisk in Table 1).

**Figure 1.**
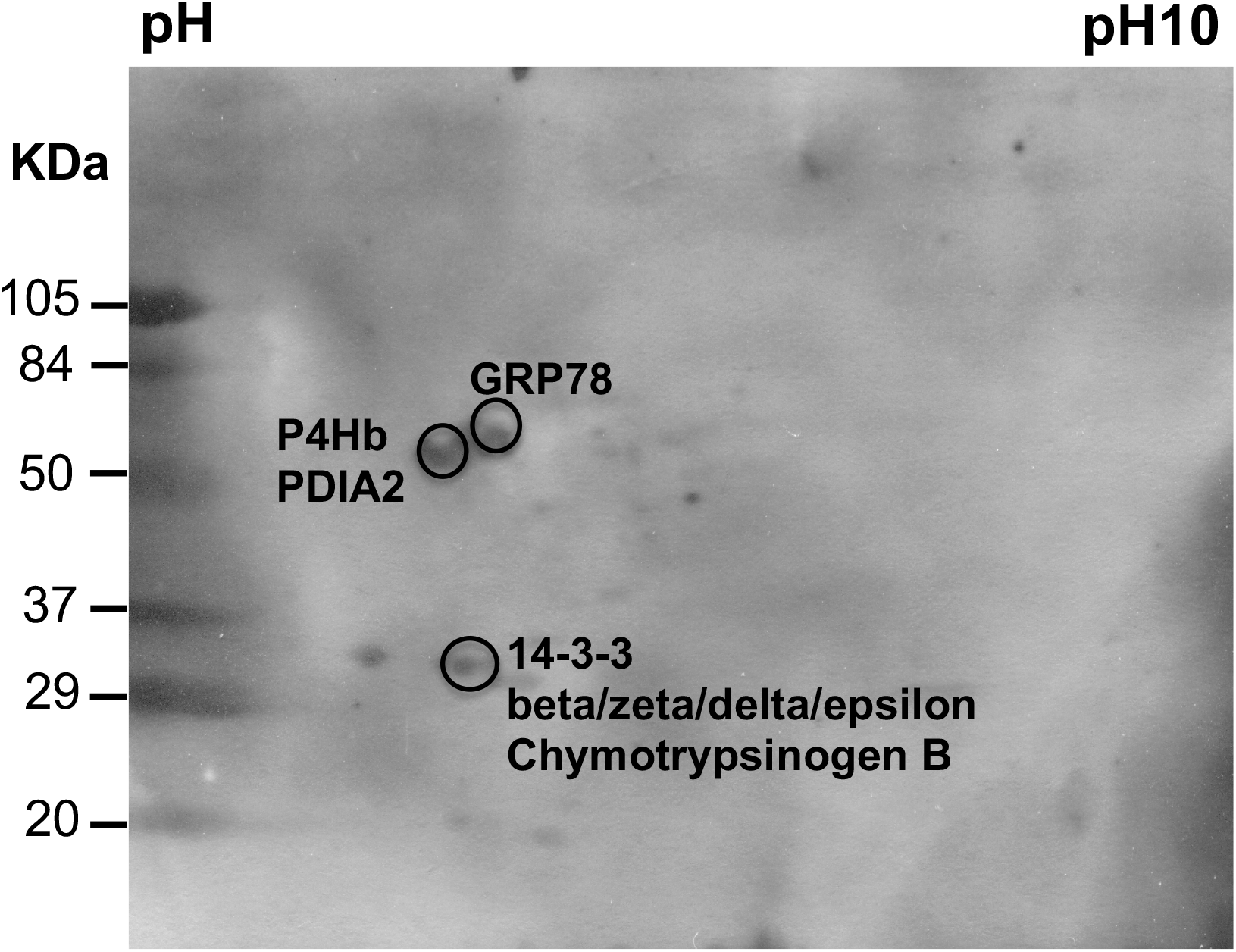
P4Hb is a major antigenic islet protein recognized by diabetic NOD serum. Two-dimensional immunoblot of 5-wk-old NOD islet extracts was probed with serum IgG from a 20-wk-old diabetic NOD mouse. The reactive islet proteins were identified by LC MS/MS spectrometry as indicated respectively. Molecular mass, in kDa, is indicated on the left.

### Oxidative and cytokine stress of human islets induces carbonyl modification of P4Hb coincident with increased proinsulin-to-insulin ratio

In human type 1 diabetes, pathology of the pancreatic islets of Langerhans develops upon exposure to proinflammatory cytokines and reactive oxygen species leading to lymphocyte infiltration. In modeling this process, we incubated normal human islets with either H_2_O_2_ or a cocktail of proinflammatory cytokines (IFNγ, IL-1β and TNFα) and assessed the induction of carbonylated P4Hb. Our results showed that carbonyl modified-P4Hb levels were significantly elevated in human islets treated with either H_2_O_2_ or a cytokine cocktail compared to untreated human islets (Fig. 2a).

**Figure 2.**
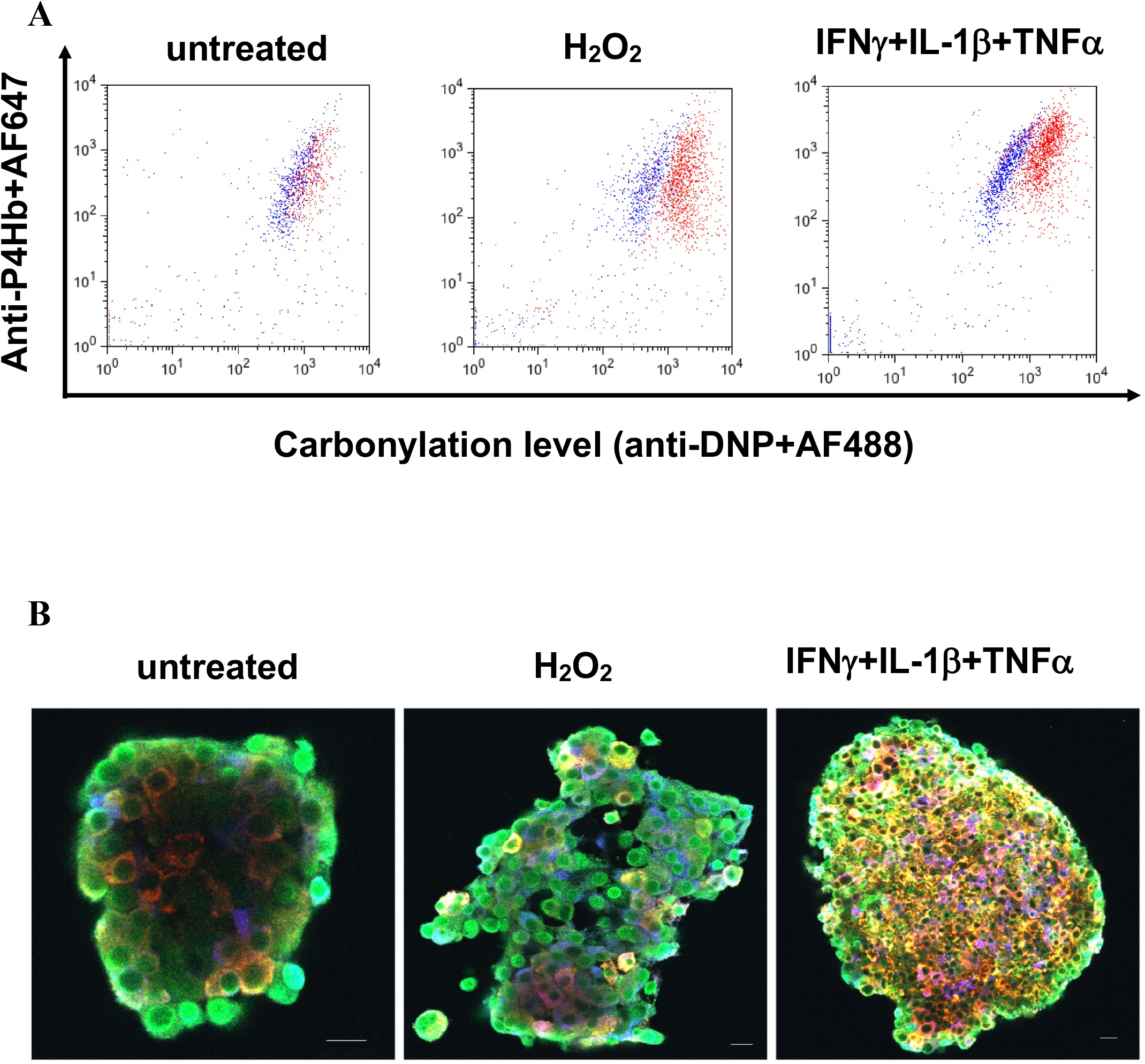
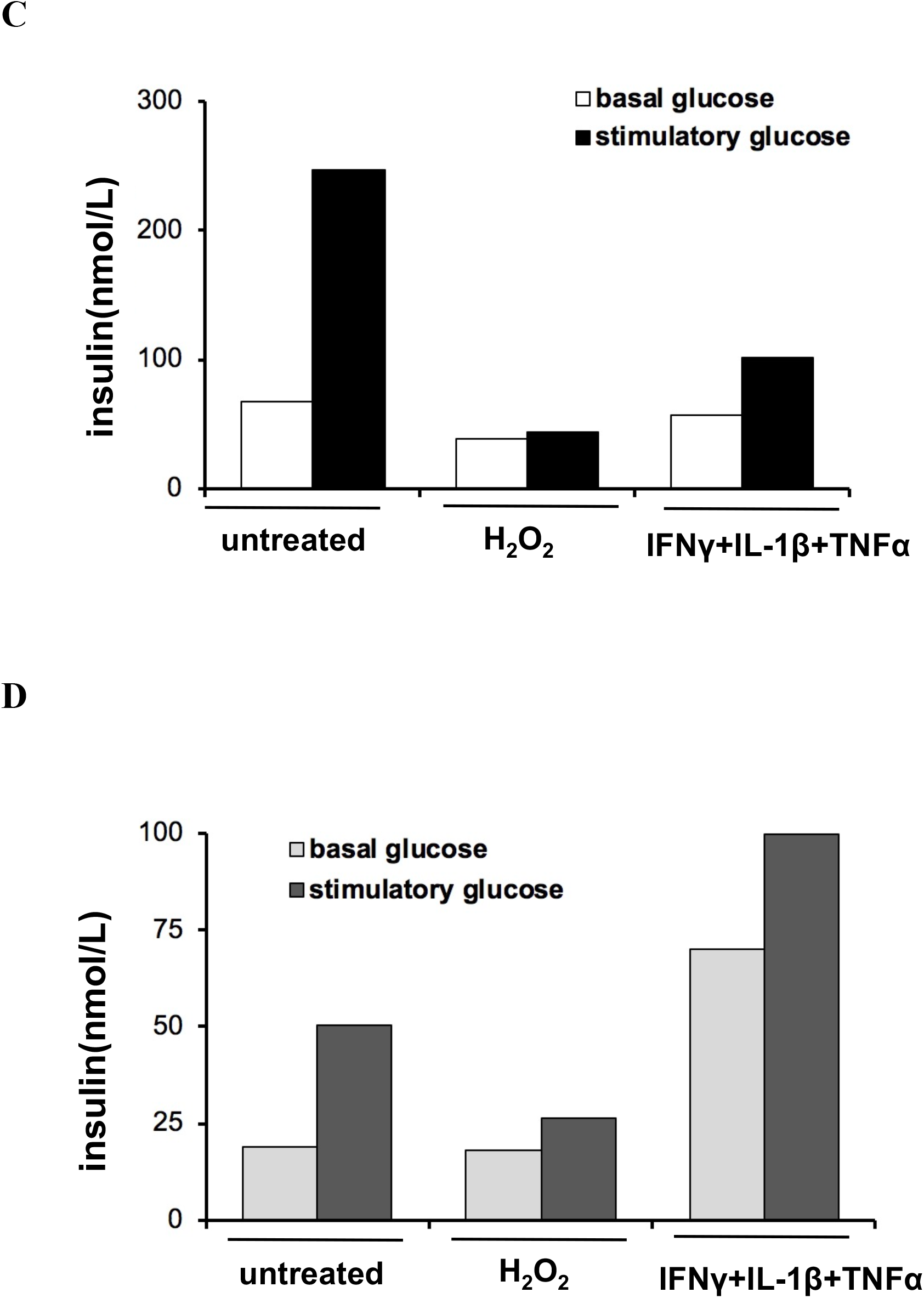
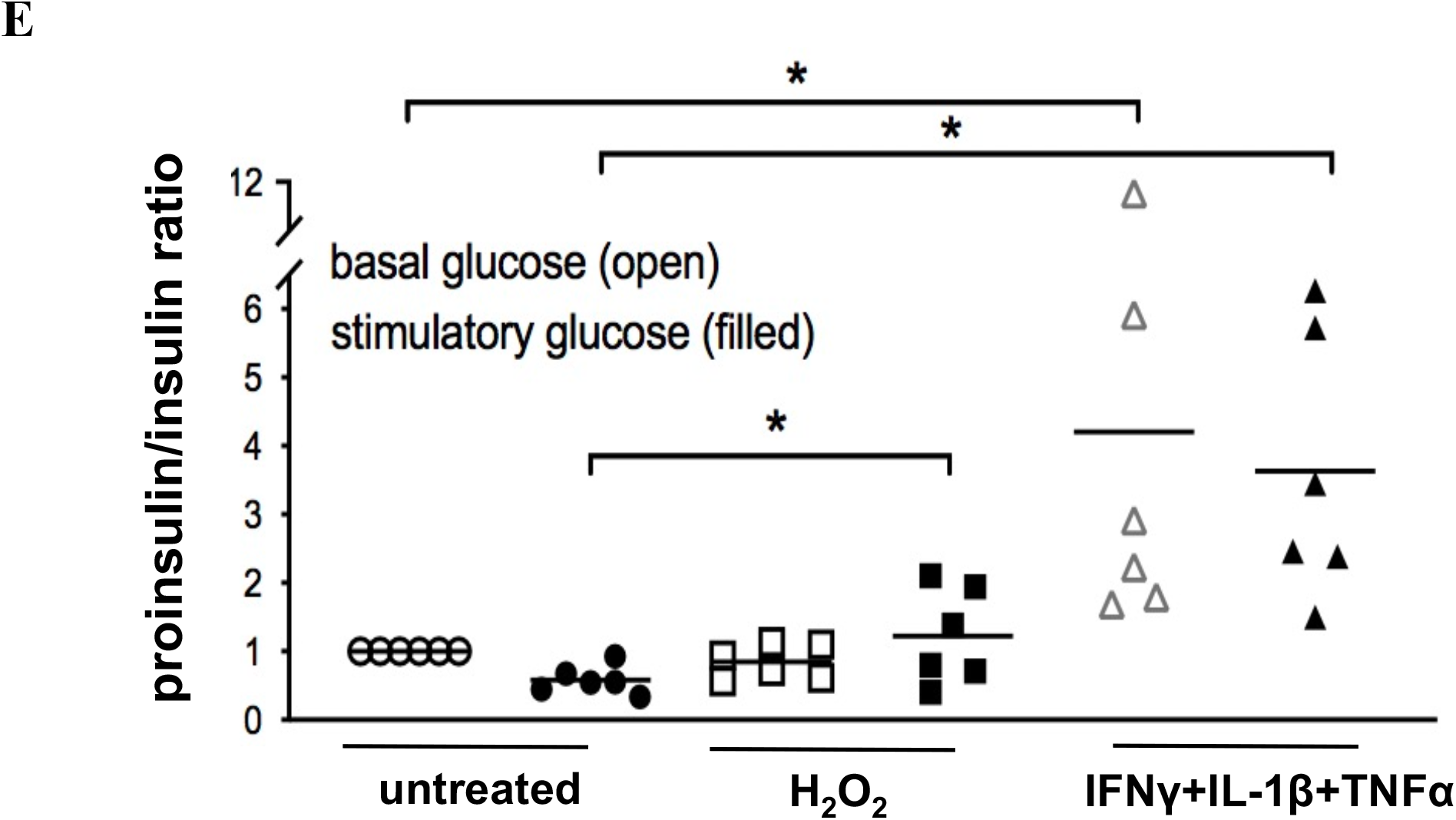
Increased carbonylation of P4Hb is coincident with impaired glucose-stimulated insulin secretion in stressed human islets. Human islet preparations were incubated in the presence or absence of oxidative stress (H_2_O_2_) or cytokine cocktail (IFNγ+IL-1β+TNFα) and followed by incubation of 5.5mM (basal) or 16.7mM (stimulatory) glucose for 1 hr. *A*: Flow cytometry analyzing the carbonylation of P4Hb in H_2_O_2_- or cytokine cocktail-treated human islets in the presence of basal glucose condition. The content of protein carbonylation is demonstrated as the AF-488 fluorescence intensity from DNPH solution-treated cells (indicated as red population) compared to the background intensity from non-DNPH treated cells (indicated as blue population) for each individual sample, respectively. *B*: After 16.7mM glucose stimulation, islets were analyzed by confocal microscopy with triple immunofluorescence for insulin (blue), P4Hb (red) and carbonylation (green). Data were shown as merged images with scale bars, 11.7µm. *C-E*: Intact human islets were incubated with basal or stimulatory glucose after treatment with H_2_O_2_ or cytokines. Supernatant was collected and secretory insulin or proinsulin was measured by ELISA. One representative experiment out of 6 human islet donor experiments is shown (*A-D*). The secretory proinsulin/insulin ratio from 6 individual stressed human islets in the presence of basal or stimulatory glucose was shown in (*E*). *, p<005.

Protein disulfide isomerase (PDI) is critical to the accurate folding of proteins within endoplasmic reticulum (ER) and cellular secretory granules, notably, the folding of insulin during its biosynthesis. In particular, P4Hb is responsible for the retention and accurate folding of proinsulin within the ER in pancreatic β cells (24). Therefore, we next determined if carbonylated-P4Hb affects metabolic functions of human β cells, including glucose-stimulated insulin secretion. The collection process of human islets does impart some cellular stress leading to carbonyl modification around the periphery of islets (green rim in Fig. 2b, left panel) as well as the presence of isoaspartyl protein modifications (data not shown). In oxidation-stressed islets, we observed that protein carbonylation is increased in β cells after glucose stimulation compared to untreated islets (increase in central green color compared to untreated islets; Fig. 2b, middle panel). Blue-green merged cells are not increased because insulin synthesis and secretion is not increased under these conditions (Fig. 2c). However, the greatest signal of carbonylated-P4Hb merged with insulin in β cells arises after glucose stimulation combined with cytokine-stressed islets (indicated as white in merged images in the right panel of Fig. 2b).

Importantly, isolated human islets have impaired glucose-stimulated insulin secretion under oxidative or cytokine stress (Fig. 2c). From normal (non-type 1 diabetes) human islets (n=6), insulin release increased an average of 3.0±1.6 fold between stimulatory (16.7mM) and basal (5.5mM) glucose exposure. Human islets were similarly treated with basal or stimulatory glucose and either H_2_O_2_ or inflammatory cytokines (IFN*γ*, IL-1*β*, and TNF*α*). Insulin release was significantly reduced in both groups, to 1.7±0.6 fold in H_2_O_2_ treated islets and to 1.3±0.8 fold in cytokine treated islets. However, the secretory proinsulin level is disproportionately elevated in cytokine treated islets compared to untreated islets either with basal or stimulatory glucose exposure (one representative islet donor is shown in Fig. 2c and 2d). Therefore, islets treated with cytokines significantly increase ratio of proinsulin over insulin concentration in secretory content compared to untreated islets (Fig. 2e). These findings demonstrate that the increase in carbonylated-P4Hb is coincident with disturbance of insulin biosynthesis and secretion in human pancreatic β cells.

### P4Hb is carbonylated under oxidative stress in beta cells and is immunoreactive to type 1 diabetes serum

Diabetic conditions lead to oxidative stress, which is known to be involved in the progression of pancreatic beta cell dysfunction. Treatment of the rat beta INS-1 cells with H_2_O_2_, the carbonylation content is increased in a dose-dependent manner (Fig. 3a) due to the intracellular reactive oxygen species (ROS) is generated by H_2_O_2_ exposure (Fig. 3b). Next, we used 300 µM H_2_O_2_-treated INS-1 cell lysate which carries a decent amount of intracellular carbonylated proteins (Fig. 3a) as a bait to ask if there is any immunoreactivity of the serum from type 1 diabetes patients. As shown in Fig. 3c, type 1 diabetes patients serum recognized one protein band which has the similar molecular weight with P4Hb (lane 1 and 3). When type 1 diabetes patient’s serum was preincubated with the recombinant human P4Hb (rhP4Hb), the band was abolished (lane 2 and 4), suggesting P4Hb contains epitopes which the autoantibody from type 1 diabetes patients reacted with.

**Figure 3:**
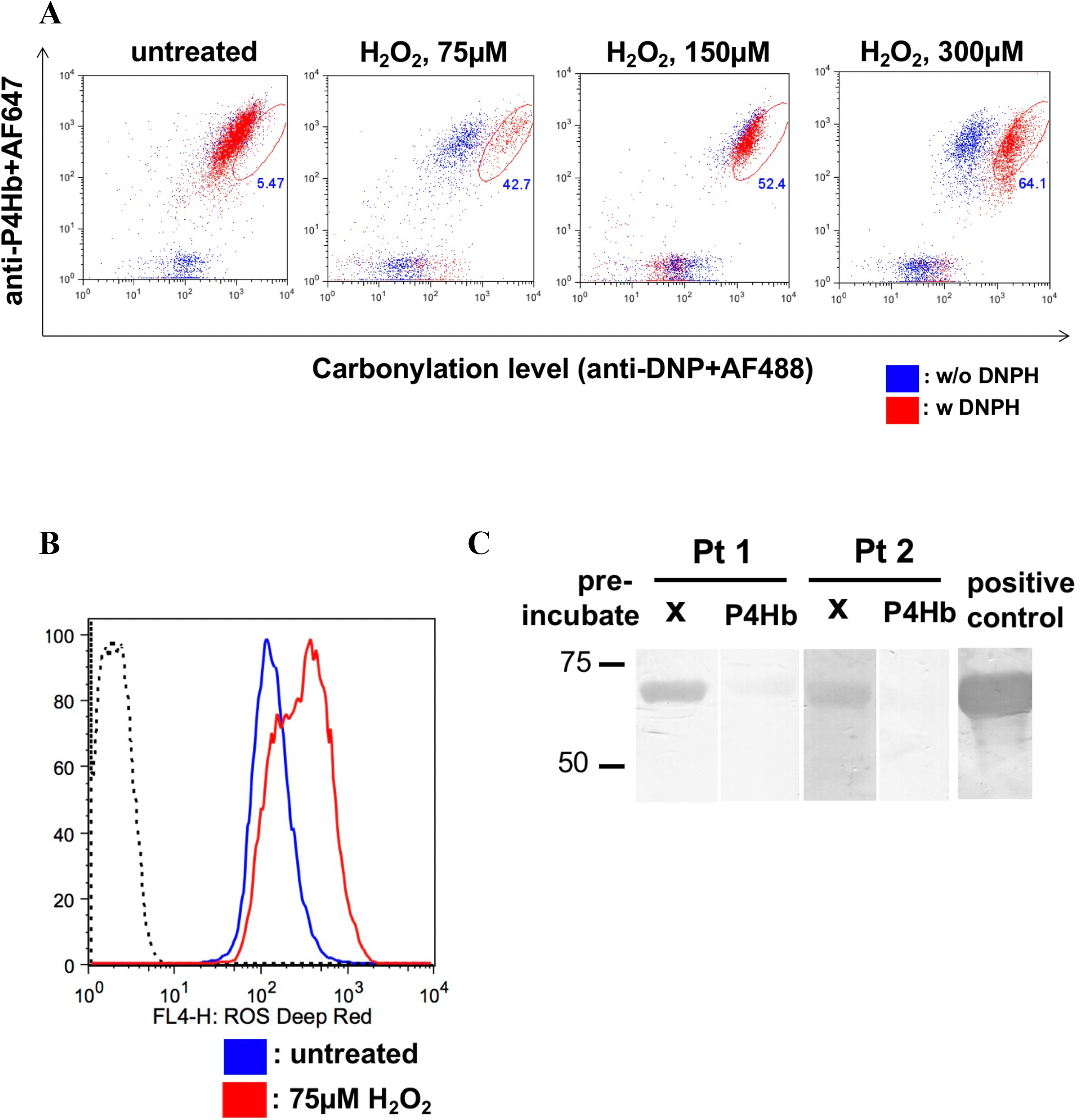
P4Hb is carbonylated under oxidative stress and is immunoreactive to type 1 diabetes serum. INS-1 cells were treated with different concentrations of H_2_O_2_ for 1 h and analyzed carbonylation by DNPH as described in Methods (*A*) and reactive oxygen species (ROS) by deep red dye (*B*) by flow cytometry, respectively. *C*: The H_2_O_2_ (300µM)-treated INS-1 lysate was electrophoresis in SDS-PAGE and transferred onto nitrocellulose membrane. Followed by immunoblot with mouse polyclonal anti-P4Hb (abcam) as positive control and sera from two established type 1 diabetes patients (Pt 1 and Pt 2). Sera were pre-incubated with rhP4Hb before immunoblot as indicated.

### NOD mice have circulating autoreactive T lymphocytes and autoantibodies that recognize P4Hb prior to hyperglycemia and islet pathology

We next evaluated cellular and humoral immune responses to P4Hb over the course of disease development in NOD murine type 1 diabetes. Isolated CD4 T cells from both pre-diabetic and diabetic NOD mice and age-matched C57Bl/6 mice were stimulated with titrating concentrations of rhP4Hb protein. While little proliferative response to rhP4Hb was observed in C57Bl/6 mice, there is a clear dose-dependent proliferative response in both prediabetic (4-5 wks of age; as shown in Fig. 4a) and diabetic (20 wks of age and blood glucose content greater than 250 mg/dl) female NOD mice. In addition, there was no difference in proliferative response to rhP4Hb between male and female NOD mice (data not shown).

**Figure 4.**
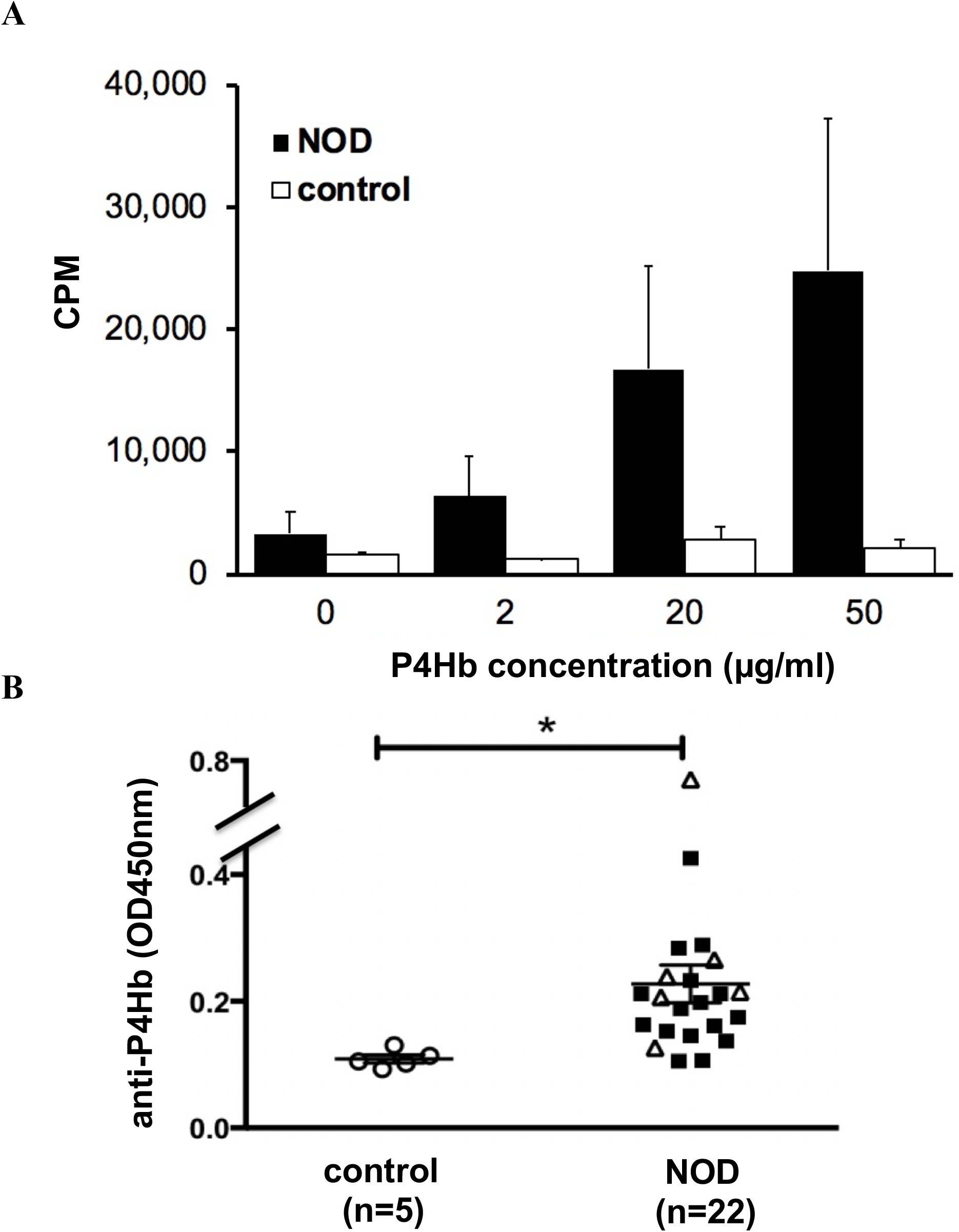
NOD mice have circulating autoreactive lymphocytes and IgG autoantibodies against P4Hb protein. *A*: Fresh isolated lymph node cells from 4- to 5-wk-old prediabetic NOD mice or control mice (C57Bl/6 strain) were incubated with varying concentrations of recombinant human P4Hb (rhP4Hb). Proliferation was measured at 72 h by [3H]thymidine incorporation. *B*: The serum levels of anti-rhP4Hb in NOD and control mice (BALB/c strain) were measured by ELISA. The number of mice analyzed is indicated at the bottom of each group. *Student t test, p<0.001. The filled squares show data where the blood glucose content was less than 250 mg/dL; the open triangles data where the blood glucose was greater than 250 mg/dL. Error bars indicate SEM of mean.

Solid phase serum immunoassays demonstrate the anti-P4Hb IgG titer was significantly higher from 4-28wk old NOD mice compared to 15-20wk old BALB/c mice (Fig. 4b). We similarly found that 20-wk-old NOR mice also had detectable IgG autoantibodies to rhP4hB (data not shown). In the related non-obese related (NOR) mouse model, there is a profound block at the peri-insulitis phase while islet-specific autoantibodies still develop. In parallel, serum samples from pre-diabetic and diabetic NOD mice and control BALB/c mice were tested for anti-insulin autoantibody levels. As shown in Table 2, all diabetic NOD mice had anti-insulin IgG antibody and 5 out of 6 mice had anti-P4Hb IgG antibody. In pre-diabetic NOD mice (n=16), 14 mice had anti-P4Hb IgG antibody while 6 mice had anti-insulin IgG antibody. Only 2 mice (of 22) lack autoantibody to either P4Hb or insulin. Data from Table 2 indicates that anti-P4Hb autoantibodies arise early in disease and precede or arise coincident the onset of autoimmunity to insulin.

**Table 2.** Autoantibodies against P4Hb are present prior to the presence of anti-insulin and hyperglycemia in NOD mice.

| **NOD mice** | **age**<br>(weeks old, wks) | **blood glucose <sup>^</sup>**<br>(blood glucose, mg/dL) | **anti-P4Hb \*** | **anti-insulin \*** |
| --- | --- | --- | --- | --- |
| #1 | 4 | — ( 123) | + | - |
| #2 | 4 | — ( 138) | + | - |
| #3 | 4 | — ( 180) | + | - |
| #4 | 4 | — ( 204) | + | - |
| #5 | 4 | — ( 88 ) | - | - |
| #6 | 4 | — ( 194) | - | - |
| #7 | 8 | — ( 104) | + | + |
| #8 | 8 | — ( 96 ) | + | + |
| #9 | 8 | — ( 93 ) | + | - |
| #10 | 8 | — ( 159) | + | - |
| #11 | 8 | — ( 116) | + | - |
| #12 | 8 | — ( 99) | + | + |
| #13 | 11 | — ( 60 ) | + | + |
| #14 | 11 | — ( 178) | + | + |
| #15 | 28 | — ( 113) | + | - |
| #16 | 28 | — ( 167) | + | + |
| #17 | 20 | + (>500) | + | + |
| #18 | 22 | + ( 466) | + | + |
| #19 | 22 | + (>500) | + | + |
| #20 | 24 | + ( 363) | + | + |
| #21 | 26 | + (>500) | - | + |
| #22 | 28 | + (>500) | + | + |

<sup>^</sup>, diabetics defined as positive if blood glucose concentration > 250 mg/dL
\*, defined as positive if ELISA OD > (average OD + 2SD) of healthy individuals (n=10)

### Prevalence of autoantibody against human P4Hb in type 1 diabetes patients

Then, we assessed P4Hb specific autoimmunity in patients with type 1 diabetes. Patients with type 1 diabetes, either early onset disease and before insulin treatment (n=21, C peptide >0.4pmol/ml) or from established type 1 diabetes (n=12, <0.3pmol/ml, from 5-40 years after type 1 diabetes diagnosis), have significantly higher anti-P4Hb IgG titer compared to healthy subjects (n=10) (Fig. 5). Moreover, 24% of early onset type 1 diabetes sera and 66% of established type 1 diabetes sera have both anti-P4Hb and anti-insulin autoantibody (Table 3). Of interest, 52% of early onset type 1 diabetes sera have anti-P4Hb autoantibody in the absence of anti-insulin antibody. Thus, the majority of early onset type 1 diabetes patients (76%) have either anti-P4Hb alone or anti-P4Hb linked with anti-insulin autoantibodies. No serum either from early onset or established type 1 diabetes had anti-insulin IgG titer without anti-P4Hb autoantibodies. Among these patients, there are eight type 1 diabetes patients who had serum collected before and after insulin therapy and the presence of anti-insulin and anti-P4Hb at different time course were shown in Table 4. Collectively, the observations in both mouse and human T1 illustrates the linked autoimmune response between P4Hb and insulin.

**Figure 5.**
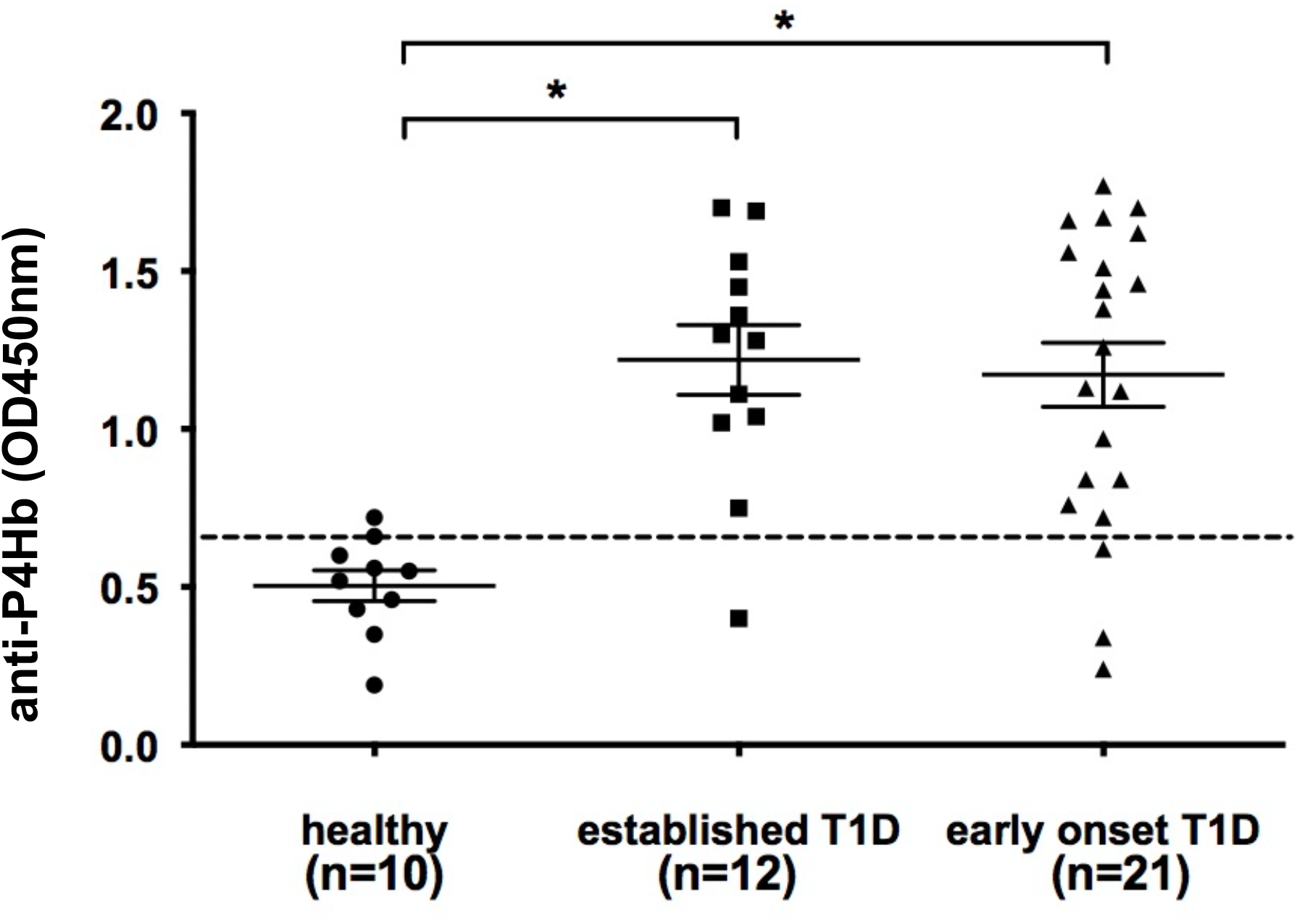
Autoantibodies against P4Hb arise in human type 1 diabetes. The serum levels of anti-rhP4Hb from early onset type 1 diabetes patients (n=21, C peptide >0.4pmol/ml) or from established type 1 diabetes (n=12, <0.3pmol/ml, from 5-40 years after type 1 diabetes diagnosis) and healthy controls (n=10) were measured by ELISA. The number of human samples analyzed is indicated at the bottom of each group and the error bars indicated SEM of mean. Cut-off line is shown as dotted line and is defined as the value of average OD_405nm_ + 2SD from healthy individuals. *Student t test, p<0.001.

**Table 3.**
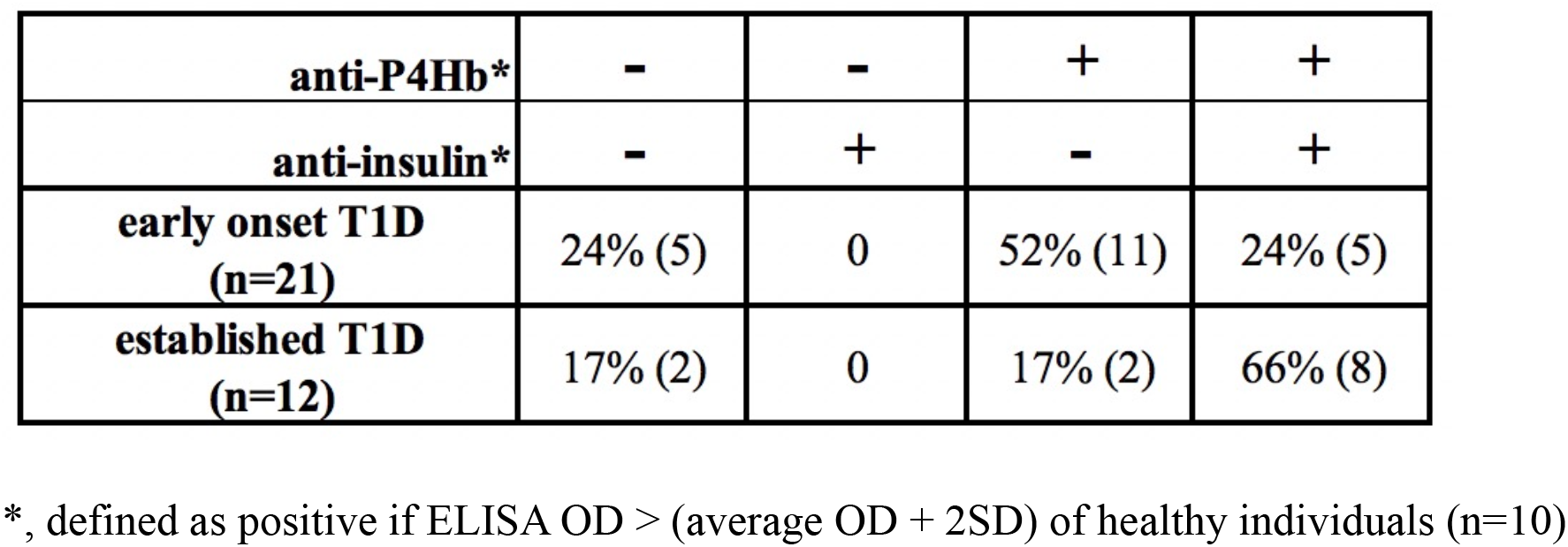
Autoantibodies against prolyl-4-hydroxylase beta peptide (P4Hb) are detected in early onset type 1 diabetes and long-established type 1 diabetes patients.

**Table 4.**
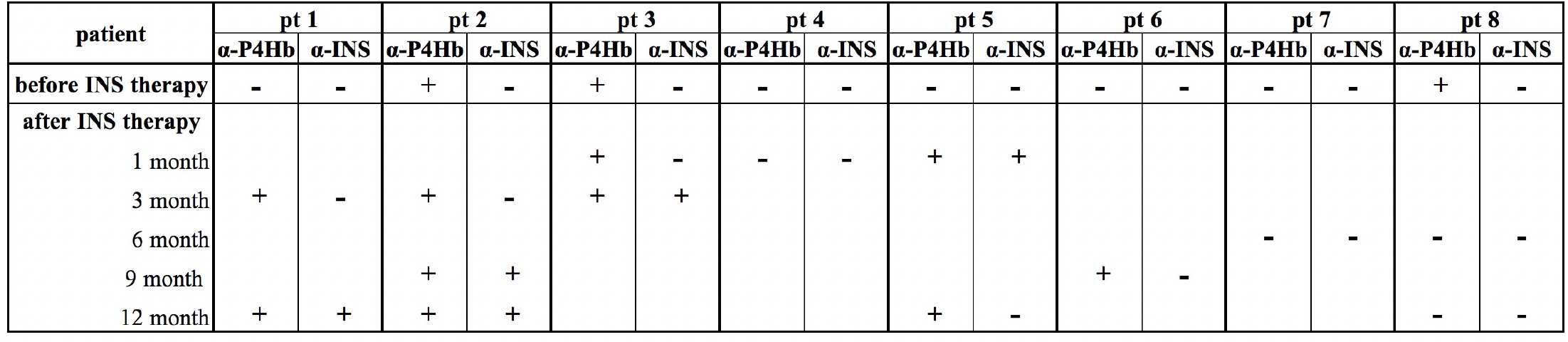
Anti-P4Hb and anti-insulin autoantibodies in serial serum samples from human type 1 diabetes.

### The identification of carbonyl residues in human P4Hb

To map the carbonylation sites of P4Hb, rhP4Hb was *in vitro* carbonylated by oxidative Fenton reaction (25). The carbonyl modification of oxidative rhP4Hb was confirmed with OxyBlot as previously described (supplementary Fig. 2a and 2b). By using computational tool, PTIK, aa 100-103, in human P4Hb protein is the only one predicted carbonylation region by Carbonylated Sites and Protein Detection (CSPD) developed by Maisonneuve *et al* (26). Interestingly, besides the Thr^101^ among in PTIK hotspot, we identified five more carbonylation sites in oxidative rhP4Hb by DNPH-assisted MS. Of note, one carbonylation site (Lys 31) in catalytic domain was identified in both control and oxidative rhP4Hb. In total, six carbonylation sites were identified in oxidative-rhP4Hb including three residues in catalytic sites and three residues in non-catalytic/substrate binding domain (Fig. 6a) and one representative MS/MS spectra of carbonyl-P4Hb peptide was shown in Fig. 6b (the detail of LC-MS/MS data is summarized in supplementary Table 1).

**Figure 6:**
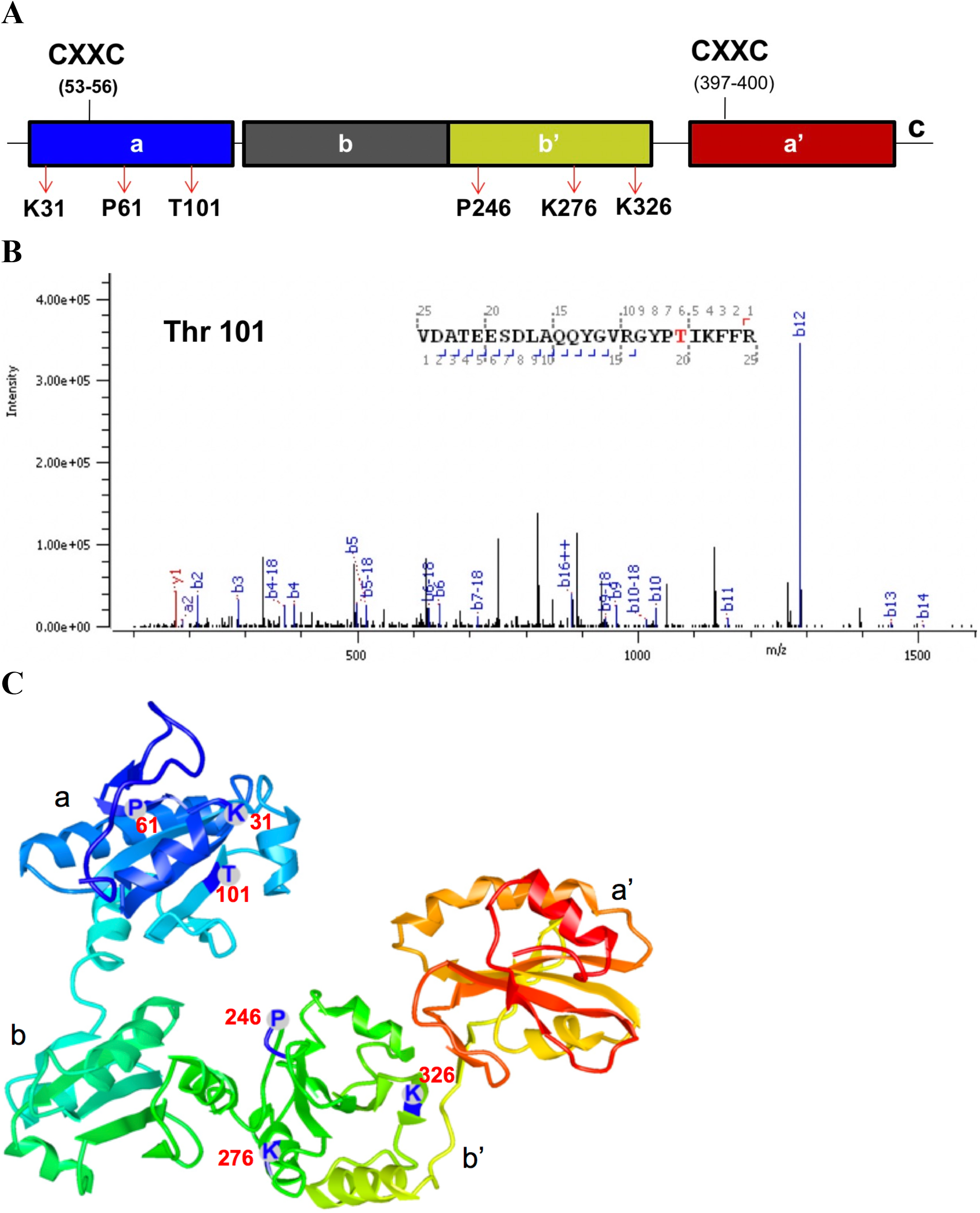
Carbonylation modifications in human P4Hb (PDB 4EKZ). *A*: The locations of carbonyl residue of human P4Hb (hP4Hb) arose during *in vitro* oxidation are indicated in the functional domains, respectively. P4Hb is composed of four thioredoxin-like domains abb’a’ and one c-terminal acidic tail, where a and a’ stand for catalytic domain with CXXC motif and b and b’ stand for non-catalytic domain. The b’ domain also contains the substrate binding site. The residual numbering is for hP4Hb with signal sequence. *B*: One representative MS/MS spectra of DNPH-derivatized peptide out of seven peptides identified in oxidative-rhP4Hb is shown. *C*: The carbonylation sites of K31, P61, T101 in domain a and P246, K276 and K326 in domain b’ are indicated in hP4Hb structure.

Human PDI is a redox-regulated chaperon responsible for catalyzing the folding of secretory proteins. The crystal structures of yeast and human PDI in different redox states showed that four thioredox domains (a, b, b’ and a’) are arranged as a horseshoe shape with the CXXC active sites, in domain a and a’, facing each other (27; 28). The b’ domain of mammalian PDI has been mapped as the primary substrate binding sites (29). Compared to the reduced state of hPDI, the oxidized hPDI results in open conformation with more extended hydrophobic areas for its substrate binding (30). The six carbonyl residues identified in this work are also highlighted in the 3D structure of human P4Hb protein (Fig. 6c).

## DISCUSSION

The overall aim of this study was to develop and utilize a broad, proteomics-based screen to identify post-translationally modified proteins and evaluate their contribution to the initiation and/or progression of type 1 diabetes. The clinical and biological relevance of this work is emphasized by other studies that demonstrate critical protein modifications in the induction and diagnosis of autoimmunity (31; 32). For example, citrullination is a product of the deamination of arginine residues and a hallmark PTM in diagnostics and in the severity of rheumatoid arthritis (RA) (33). Autoantibodies against citrullinated self-proteins appear early in the disease, precede the expression of pathology in RA, and are not found in otherwise healthy individuals (34). Moreover, protein citrullination and antibodies against citrullinated self-proteins have also played an important role in type 1 diabetes (35).

Type 1 diabetes pathogenesis is linked to the generation of tissue free radicals (36). Carbonylation is the major PTM product in tissues/cells under oxidative stress. Oxidation-triggered PTMs result in increased carbonyl-modified plasma proteins in patients with type 1 diabetes (37). Moreover, hyperglycemia amplifies oxidative stress, leading to pancreatic beta cell and endothelial cell dysfunction (38). The deficiency of glutathione peroxidase 1 (GPx1), the one of the major antioxidant enzymes, leads to metabolic changes similar to type 1 diabetes and carbonylation results in enzymatically inactivation of GPx1, which might contribute to insulin resistance (39).

One novel autoantigen identified in this study, P4Hb (also known as PDIA1), is a member of the PDI family and is the beta subunit of a tetramer of prolyl-4-hydroxylase (P4H). This tetramer holoenzyme has several important cellular functions, including the hydroxylation of proline residues in procollagen. Relevant to the biology of type 1 diabetes, P4Hb activity in β cells modulates glucose-stimulated release of insulin (24). P4Hb is both a chaperone and thioreductase, participating in disulfide bond formation and isomerization during the process of insulin biosynthesis and secretion. It has been demonstrated that chemical modification of PDI, ablating the thioredoxin activity, prevents the refolding of denatured and reduced proinsulin *in vitro* (40). Moreover, proinsulin forms aggregate in the absence of the chaperone activity of the PDI. Our data is consistent with recent studies illustrating that pancreatic beta cells fail to properly process proinsulin (41).

The significance of P4Hb-associated insulin misfolding is illustrated in studies from the Akita mouse model, where improper disulfide bond formation leads to the retention of misfolded proinsulin in the β-cell ER (42; 43). Others have proposed that restoration of the unfolded protein response in pancreatic β cells protects mice against type 1 diabetes (44). Moreover, GRP78/BiP, one of the carbonyl modified antigenic islet proteins identified in this study, is also an ER chaperone. The transcriptional activity of GRP78/BiP promoter is up-regulated in response to unfolded proteins in the ER (45). Of interest, the mispairing of disulfide bonds in proinsulin increases ER stress response resulting in GRP78/BiP promoter activation in INS-1 β cells (46). Herein, our data demonstrated that carbonylation of P4Hb is greatly increased in human islets under oxidative or inflammatory cytokine stress. Moreover, insulin production is significantly reduced in stressed human islets compared to untreated human islets, coincident with carbonyl modification of P4Hb. Recently, antibodies against oxidized insulin were detected in patients with type 1 diabetes (47). P4Hb, but also insulin itself, may undergo carbonylation in pancreatic islets during insulitis and results in misfolded proinsulin or insulin, a potential initiating event leading to the onset of type 1 diabetes. this pathway explains observations of increasing proinsulin/insulin ratios in the progression of type 1 diabetes as recently reported (11; 41).

Carbonylation is an irreversible and non-enzymatic addition of aldehyde or ketone groups to specific amino acid residues including direct oxidation (Lys, Arg, Pro, Thr) and indirect oxidation (Glu, His, Cys, Trp) (7). It is not known if the oxidative carbonylation of P4Hb occurs randomly at sites along the protein or if “hot spots” exist. In our study, we found that autoantibody recognizing P4Hb was detected in prediabetic nod animals before the onset of type 1 diabetes hyperglycemia and anti-insulin autoantibody production. In human type 1 diabetes, we also found detectable autoantibodies against p4hb in patients with early-onset as well as established type 1 diabetes.

One other type 1 diabetes-associated autoantigen identified in our work, 14-3-3 family proteins, consists of at least seven isoforms. The 14-3-3 proteins have been identified as regulatory elements in insulin resistance, signal transduction and β cells survival (48; 49). In addition, 14-3-3 protein epsilon has been shown to bind to the glial fibrillary acidic protein (GFAP), an autoantigen found in type 1 diabetes and multiple sclerosis (50). In a manner similar to P4Hb, we found that sera from early-onset type 1 diabetes patients have an IgM response to recombinant 14-3-3 zeta and 14-3-3 epsilon isoforms (unpublished data).

In conclusion, we identified selected inflammation-induced carbonyl protein modifications from both mouse and human pancreatic islets. These modifications may lead to a specific loss of both B and T cell immune tolerance. This approach successfully identified both novel and known type 1 diabetes autoantigens such as GRP78 and pancreatic amylase. Anti-P4Hb autoimmunity always precedes anti-insulin autoantibodies and β-cell dysfunction in type 1 diabetes. This autoimmune pathway is marked by impaired glucose-stimulated insulin secretion and provides explanation for increased proinsulin/insulin ratios found during the progression of human type 1 diabetes. Beyond identifying novel autoimmune targets of early type 1 diabetes and altered mechanisms of insulin metabolism, these studies also provide potential antioxidant therapeutic strategies to reverse disease-relevant biochemical pathways.

## Acknowledgements

We are grateful to Professor Colin Thorpe (University of Delaware, Newark, Delaware) for providing human hPDI-pTrcHisA clone for these studies. We thank members of the Mamula laboratory, Cameron Gesick, Elaina Bruck, Emily Schroeder and Tyler Masters for excellent technical assistant for this study. We thank Ningwen Tai for technical assistant in isolation of pancreatic islets and advice for some experiments. We are grateful to Lucy Zhang for careful management of the NOD mouse colony. We thank Drs. of Carla Greenbaum and Jane Buckner for collecting and archiving human samples for this work. The authors would like to thank Navin Rauniyar, W. M. Keck Foundation Biotechnology Resource Laboratory, Yale University, for the expertise of high-resolution mass spectrometry.

## Funding

This research was supported by the Training Program in Investigative Rheumatology (5T32-AR007107) to S.E.C., Innovative Grant from the Juvenile Diabetes Research Foundation to M.J.M., and National Institutes of Health Grant (DK104205-01) to K.H. and M.J.M. L.W. was supported by National Institutes of Health Grant (DK092882) and Diabetes Research Center (DK-11-015).

## Author Contributions

M.L.Y. designed and performed the experimentation, data analysis, and composition of the manuscript. S.E.C. and R.G. performed the experimentation of 2D-gel electrophoresis and confocal microscopy, respectively. M.J.M., and K.H., directed the project and edited the manuscript. K. H. and C.S. provided key type 1 diabetes serum sets to the study. L.W. provided the NOD mice utilized in the study, supervised pancreatic islet isolation and also provided helpful discussion and manuscript edits. T.L. and J.K. performed LC MS/MS and analyzed high-resolution mass spectrometry data. E.A.J., C.S., C.E.M., S.C. and C.C. provided helpful discussion and manuscript edits.

## Conflict of Interest statement

The authors declare no competing financial interests.

## Prior Presentation

Parts of this study were presented in abstract form at the 100^th^ annual meeting of American Association of Immunologists (AAI), IMMUNOLOGY 2016^TM^, Seattle, WA, 13-17 May 2016 and the 5^th^ European Congress of Immunology (ECI) meeting, Amsterdam, 2-5, September 2018.

## Supplemental Material

**Supplemental Table 1.**
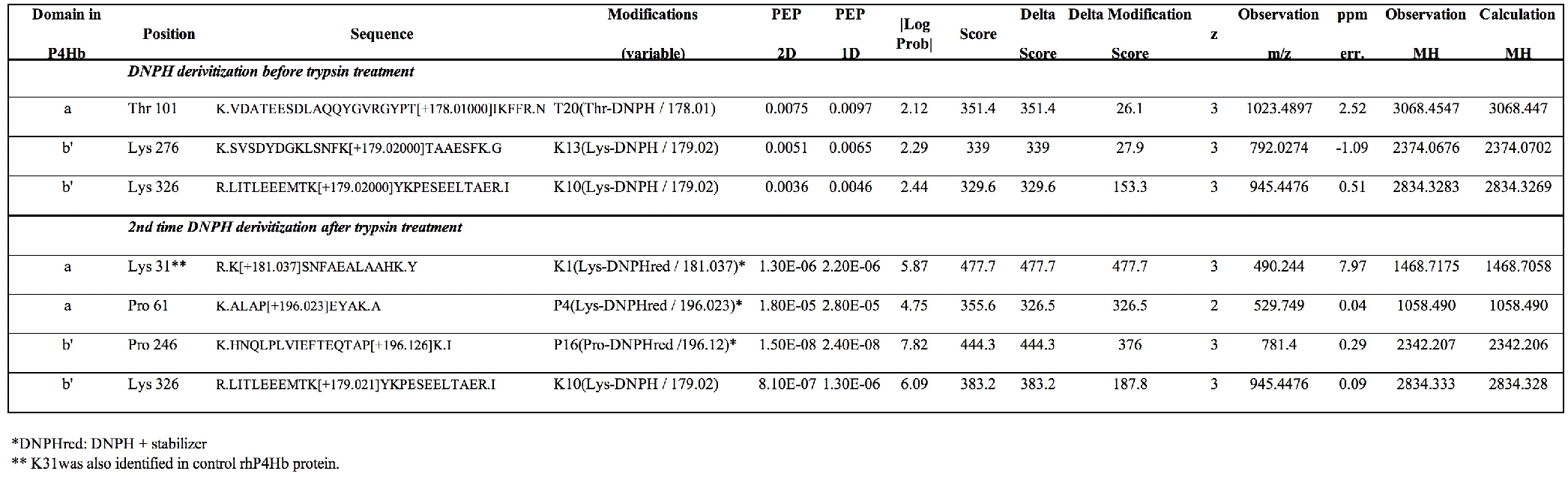
The carbonyl residues identified in human P4Hb under oxidative stress.

**Supplemental Figure 1.**
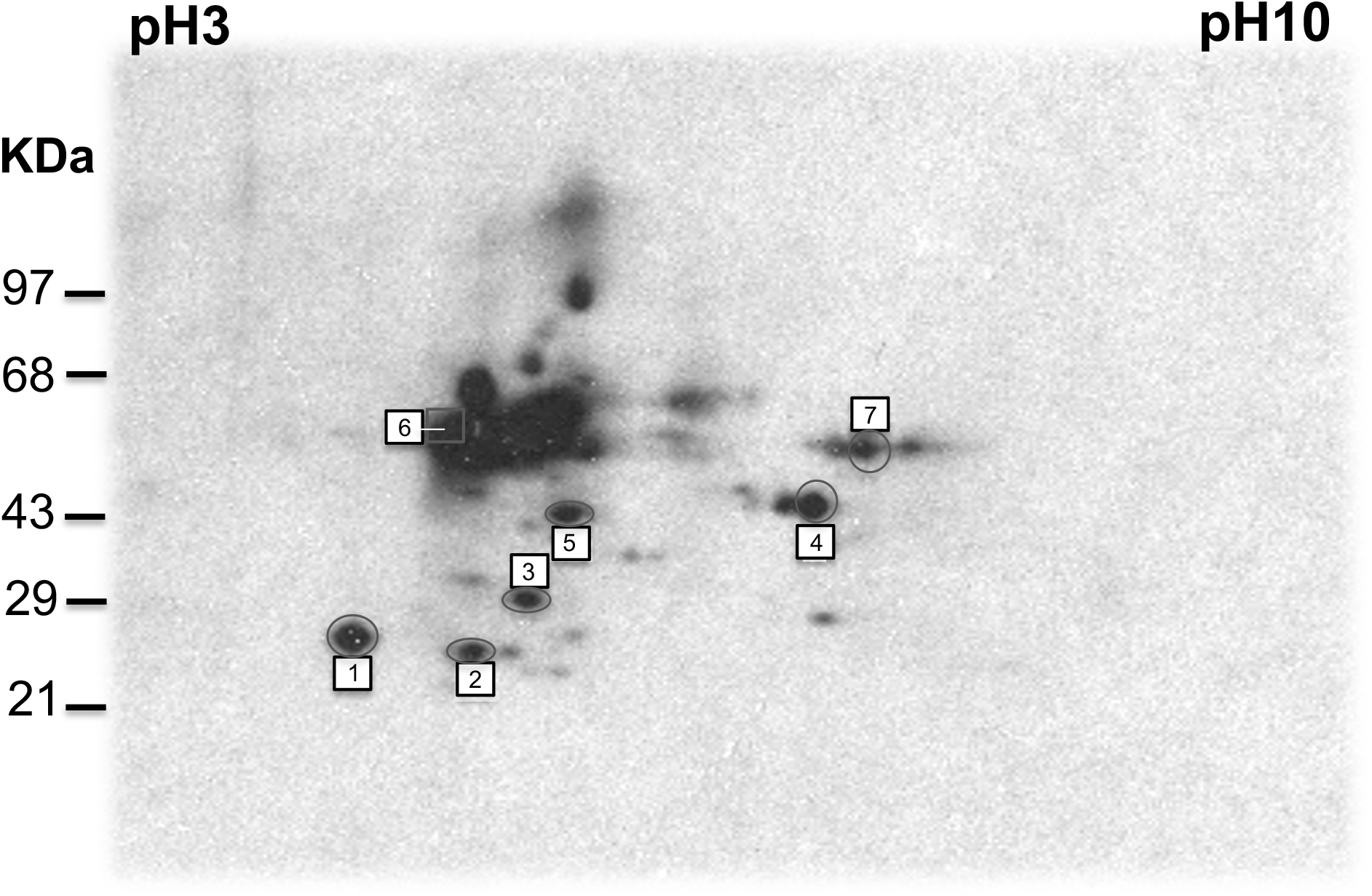
NOD islet proteomic analysis for carbonyl-modified proteins. Representative two-dimensional blot of carbonyl-modified proteins from prediabetic NOD mice. Islet proteins from 5-wk-old NOD mice were separated by IEF and derivatized with DNPH. Second dimension separation was performed in 12.5% SDS-PAGE gels. Squares/circles and numbers indicate spots identified by LC MS/MS spectrometry. Molecular mass, in kDa, is indicated on the left.

**Supplemental Figure 2:**
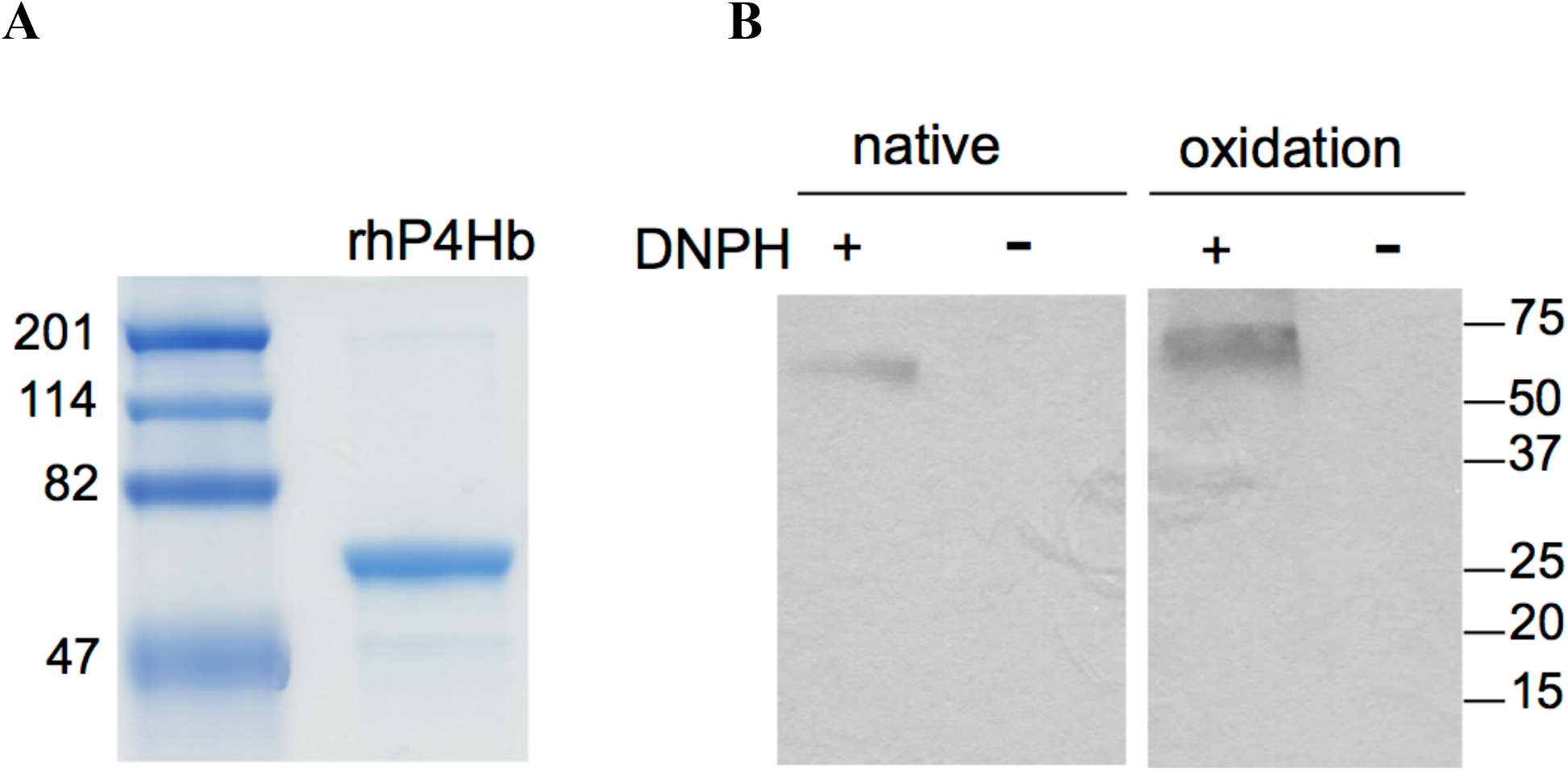
The characterization of purified recombinant human P4Hb from hPDI-pTrcHisA clone. (A) Briefly, the hPDI-pTrcHisA plasmid was transformed into BL21(DE3) cells for expression under 0.5mM IPTG induction and purified with imidazole by using Pro Bond Ni-NTA resin. Proteins were concentrated with Centriprep YM-30 and judged to be ∼95% pure by SDS-PAGE stained with Coomassie Brilliant Blue. (B) The purified rhP4Hb was incubated in PBS (native) or PBS containing 100µM FeSO4, ascorbate, 25mM H_2_O_2_ and 25mM ascorbate (oxidation) at 37°C for 4h. Then the carbonyl modification was analyzed by OxyBlot. Molecular weight marker (kDa), is indicated on the edge of the gel, respectively.

## Notes

### Competing Interest Statement

The authors have declared no competing interest.

