## Supplemental Figures for "Carbonyl Post-Translational Modification Associated with Early Onset Type 1 Diabetes Autoimmunity"

### Supplemental Material

### Supplemental Table 1.

#### The carbonyl residues identified in human P4Hb under oxidative stress.

| Domain in P4Hb | Position | Sequence | Modifications (variable) | PEP 2D | PEP 1D | \|Log Prob\| | Score | Delta Score | Delta Modification Score | z | Observation m/z | ppm err. | Observation MH | Calculation MH |
|---|---|---|---|---|---|---|---|---|---|---|---|---|---|---|
| | *DNPH derivitization before trypsin treatment* | | | | | | | | | | | | | |
| a | Thr 101 | K.VDATEESDLAQQYGVRGYPT[+178.01000]IKFFR.N | T20(Thr-DNPH / 178.01) | 0.0075 | 0.0097 | 2.12 | 351.4 | 351.4 | 26.1 | 3 | 1023.4897 | 2.52 | 3068.4547 | 3068.447 |
| b' | Lys 276 | K.SVSDYDGKLSNFK[+179.02000]TAAESFK.G | K13(Lys-DNPH / 179.02) | 0.0051 | 0.0065 | 2.29 | 339 | 339 | 27.9 | 3 | 792.0274 | -1.09 | 2374.0676 | 2374.0702 |
| b' | Lys 326 | R.LITLEEEMTK[+179.02000]YKPESEELTAER.I | K10(Lys-DNPH / 179.02) | 0.0036 | 0.0046 | 2.44 | 329.6 | 329.6 | 153.3 | 3 | 945.4476 | 0.51 | 2834.3283 | 2834.3269 |
| | *2nd time DNPH derivitization after trypsin treatment* | | | | | | | | | | | | | |
| a | Lys 31** | R.K[+181.037]SNFAEALAAHK.Y | K1(Lys-DNPHred / 181.037)* | 1.30E-06 | 2.20E-06 | 5.87 | 477.7 | 477.7 | 477.7 | 3 | 490.244 | 7.97 | 1468.7175 | 1468.7058 |
| a | Pro 61 | K.ALAP[+196.023]EYAK.A | P4(Lys-DNPHred / 196.023)* | 1.80E-05 | 2.80E-05 | 4.75 | 355.6 | 326.5 | 326.5 | 2 | 529.749 | 0.04 | 1058.490 | 1058.490 |
| b' | Pro 246 | K.HNQLPLVIEFTEQTAP[+196.126]K.I | P16(Pro-DNPHred /196.12)* | 1.50E-08 | 2.40E-08 | 7.82 | 444.3 | 444.3 | 376 | 3 | 781.4 | 0.29 | 2342.207 | 2342.206 |
| b' | Lys 326 | R.LITLEEEMTK[+179.021]YKPESEELTAER.I | K10(Lys-DNPH / 179.02) | 8.10E-07 | 1.30E-06 | 6.09 | 383.2 | 383.2 | 187.8 | 3 | 945.4476 | 0.09 | 2834.333 | 2834.328 |

*DNPHred: DNPH + stabilizer
** K31was also identified in control rhP4Hb protein.

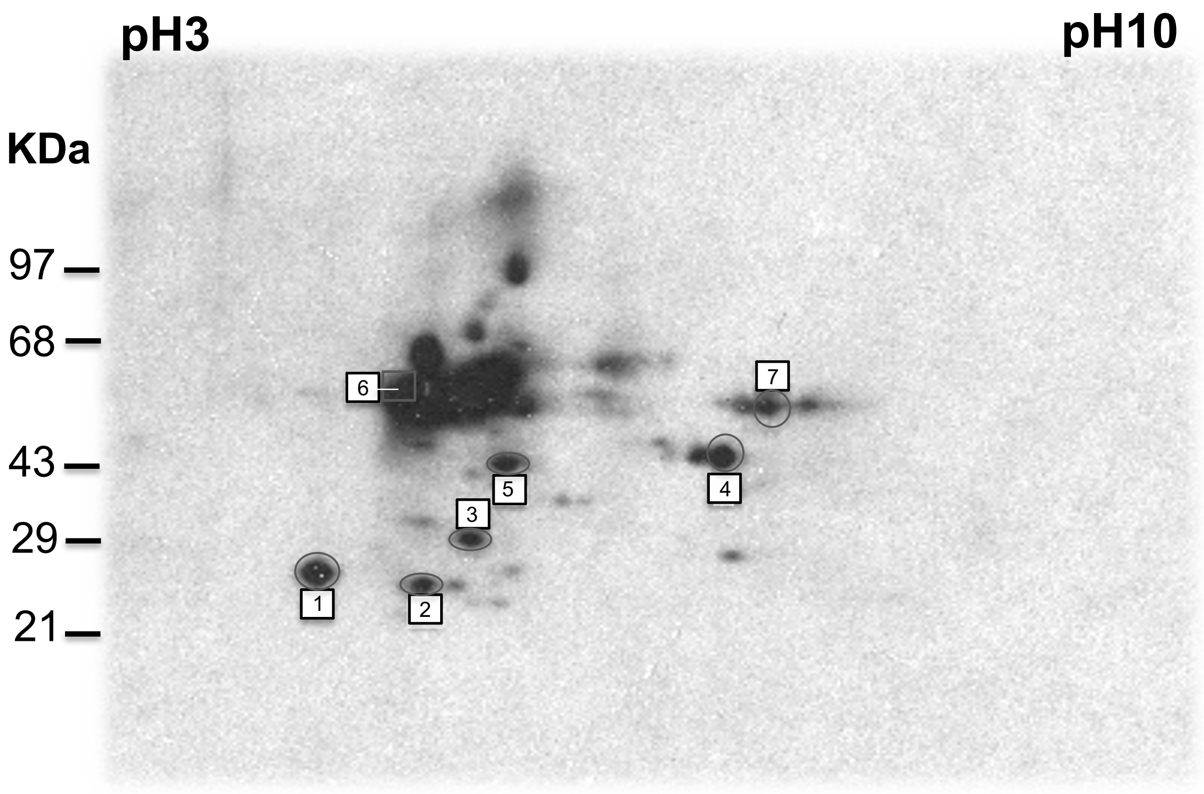


**Supplemental Figure 1. NOD islet proteomic analysis for carbonyl-modified proteins.** Representative two-dimensional blot of carbonyl-modified proteins from prediabetic NOD mice. Islet proteins from 5-wk-old NOD mice were separated by IEF and derivatized with DNPH. Second dimension separation was performed in 12.5% SDS-PAGE gels. Squares/circles and numbers indicate spots identified by LC MS/MS spectrometry. Molecular mass, in kDa, is indicated on the left.

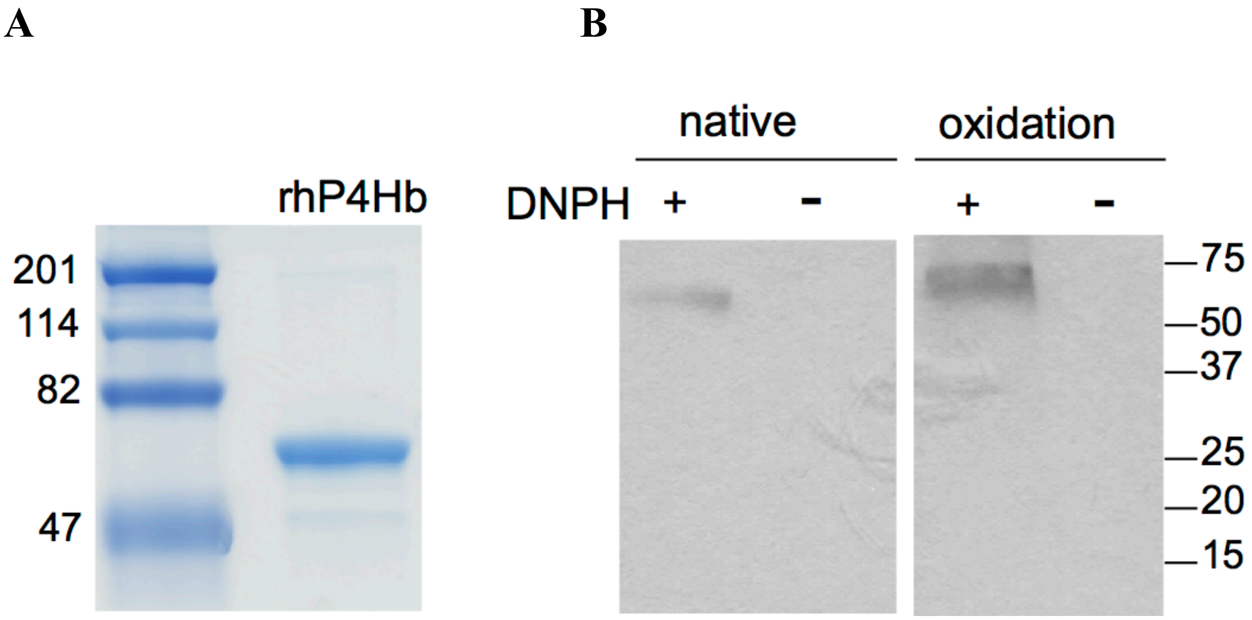


**Supplemental Figure 2: The characterization of purified recombinant human P4Hb from hPDI-pTrcHisA clone.** (A) Briefly, the hPDI-pTrcHisA plasmid was transformed into BL21(DE3) cells for expression under 0.5mM IPTG induction and purified with imidazole by using Pro Bond Ni-NTA resin. Proteins were concentrated with Centriprep YM-30 and judged to be ~95% pure by SDS-PAGE stained with Coomassie Brilliant Blue. (B) The purified rhP4Hb was incubated in PBS (native) or PBS containing 100μM FeSO4, ascorbate, 25mM $H_2O_2$ and 25mM ascorbate (oxidation) at 37°C for 4h. Then the carbonyl modification was analyzed by OxyBlot. Molecular weight marker (kDa), is indicated on the edge of the gel, respectively.
